# Episodic and semantic memory contributions to imagination and creativity

**DOI:** 10.64898/2026.02.08.704672

**Authors:** Preston P. Thakral, Kevin P. Madore, Ruth E. Gomez, Aleea L. Devitt

## Abstract

The ability to generate novel creative ideas (divergent thinking) is closely linked with our ability to imagine novel future events (episodic simulation). Here, we employed an individual differences approach to examine whether divergent thinking and episodic simulation are differentially associated with episodic and semantic retrieval ability. In response to object word cues, participants generated meanings and definitions (semantic memory), remembered a past event (episodic memory), imagined a novel future event (episodic simulation), or generated novel uses (divergent thinking). Replicating previous findings, divergent thinking ability was predicted by the number of episodic details generated during episodic simulation. When directly comparing episodic and semantic memory, the strongest predictor of divergent thinking was semantic memory. In contrast, episodic simulation ability was predicted by both episodic and semantic memory. We interpret these findings as support for the semantic scaffold hypothesis of imagination, according to which semantic memory provides the necessary scaffold or framework for flexible expressions of cognition such as divergent thinking and episodic simulation. As episodic simulation, relative to divergent thinking, was associated with both episodic and semantic retrieval, these findings are taken to reflect common reliance on event construction processes recruited during both episodic remembering and imagining.

---

Research on creativity has focused on two distinct forms of creative thinking; convergent and divergent thinking. Convergent thinking refers to the ability to generate the single best solution to a problem. In contrast, divergent thinking refers to the ability to combine diverse bits information and generate as many novel solutions to a given problem as possible. Both forms of creative cognition have long been considered to rely on the flexible association of concepts stored in semantic memory (Mednick, 1962). For example, prominent models of creativity emphasize that creative thinking is associated with greater conceptual expansion (e.g., Abraham et al., 2014), executive control over the variability in semantic network activation (e.g., Hills, & Kenett, 2025), more random sampling/searching of semantic memory (e.g., Hass, 2017), and less modularity and greater flexibility in semantic memory network organization (e.g., Kenett, Anaki, & Faust, 2014; Kenett, Levy, Kenett, Stanley, Faust, & Hvalin, 2018; for reviews, see Abraham & Bubic, 2015; Beaty & Kenett, 2023; Benedek, Beaty, Schacter, & Kenett, 2023; Kenett, 2025).

While both divergent and convergent creativity are linked to semantic processing (for a review, see Gerver, Griffin, Dennis, & Beaty, 2023), there is now a wealth of evidence to indicate that divergent relative to convergent thinking draws on episodic memory as well (i.e., the ability to recollect a specific past episode with detailed who, what, where, and when information; for a review, see Ditta & Storm, 2018; Benedek et al., 2023). For example, participants report thinking of past events when generating creative ideas in the Alternate Uses Task (AUT), a laboratory measure of divergent thinking where participants generate unusual and creative uses for an everyday object (Gilhooly, Fioratou, Anthony, & Wynn, 2007). Patients with damage to the hippocampus who show deficits in episodic memory also show deficits in verbal tests of divergent thinking, like the AUT, and visual tests of divergent thinking, like the figural form of the Torrance Tests of Creative Thinking (Duff, Kurczek, Rubin, Cohen, & Tranel, 2013).

Moreover, studies have shown that after receiving a brief training in episodic retrieval (i.e., an episodic specific induction), participants generate more uses in the AUT relative to a training focused on non-episodic retrieval (Madore, Addis, & Schacter, 2015; Madore, Jing, & Schacter, 2016; Madore, Thakral, Beaty, Addis, & Schacter, 2019).

There is likewise evidence from neuroimaging studies indicating neural overlap during episodic memory and divergent thinking. For example, in Beaty, Thakral, Madore, Benedek, and Schacter (2018), relative to a non-episodic/semantic control task, common neural activity during both episodic memory and divergent thinking was observed in the ‘core network’, a set of regions which includes the hippocampus that has been consistently associated with episodic event construction (Benoit & Schacter, 2015).

Linked to both divergent thinking and episodic memory is episodic simulation: the construction of a detailed mental representation of a specific and novel autobiographical future event (Szpunar, Spreng, & Schacter, 2014). It is now widely recognized that episodic simulation, akin to divergent thinking, draws heavily on episodic retrieval, in that it requires the retrieval and recombination of specific autobiographical information (e.g., Moscovitch, Cabeza, Winocur, & Nadel, 2016; Schacter & Addis, 2007, 2020; Schacter & Thakral, in press; Sheldon & Levine, 2016). According to the memory in creative ideation (MemiC) framework (Benedek et al., 2023), episodic simulation is a key component process to successful divergent thinking. In the MemiC framework, creatives ideas arise from four processing stages. The first two stages involve individuals searching and retrieving episodic or semantic information, and then constructing ideas based on that retrieved information. The final two stages involve the evaluation of the generated idea. It is during this stage where simulation plays a key role: individuals run an episodic simulation to assess the ideas’ properties and feasibility (i.e., novelty and effectiveness). If the idea is judged as neither novel nor effective, the generation process is reengaged. A similar theoretical proposal was raised by Roberts and Addis (2018), whereby divergent thinking is considered a form of simulation analogous to imagining future events, as they both rely on the ability to form associations between retrieved episodic and semantic information (among other shared processes such as executive functions that act on those associations).

Recent research indicates that, like divergent thinking, episodic simulation is supported by semantic retrieval (for reviews, see Schacter et al., 2012; Irish & Piguet, 2013; Klein, 2013; Irish & Piolino, 2016; Szpunar, Spreng, & Schacter, 2014; Macleod, 2016; Roberts & Addis, 2018). In one study, measures of executive function, which included semantic fluency, correlated with the specificity of future but not past events (D’Argembeau, Ortoleva, Jumentier, & van der Linden, 2010). Other behavioral work has shown that verbal descriptions of participants imagined future events contain greater amounts of autobiographical knowledge than remembered past events (e.g., personal semantic information; D’Argembeau & Mathy, 2011). Related evidence comes from patients diagnosed with semantic dementia resulting from damage to the anterior temporal lobes, who exhibit deficits in semantic memory. These patients also demonstrate a selective reduction in the generation of episodic details when imagining novel future events, but not when remembering past events (Irish, Addis, Hodges, & Piguet, 2012a; Irish, Addis, Hodges, & Piguet, 2012b; Irish et al., 2016; see also, Klein, Loftus, & Kilhstrom, 2002). Complementing these studies, Duval et al., (2012) showed that semantic dementia patients demonstrate reduced self-projection abilities to the future but not past, as well as a reduced sense of pre-but not re-living (i.e., autonoetic consciousness). Further consistent with this patient data are neuroimaging studies that have revealed greater activity in brain regions associated with semantic memory (e.g., lateral temporal cortex) during imagining the future relative to remembering the past (e.g., Addis, Pan, Vu, Laiser, & Schacter, 2009; Addis, Roberts, & Schacter, 2011).

These links between future simulation and semantic memory are theorized to reflect the role of semantic memory in serving as a ‘scaffold’ or framework within which novel future events can be constructed (e.g., Irish et al., 2012a). According to this view, semantic memory provides the necessary schema or abstracted representation for novel future event imagination, providing the conceptual framework upon which episodic details (e.g., who, what, where, and when) can be integrated to generate a novel and coherent future event The evidence reviewed above suggests that episodic simulation and divergent thinking are associated with the retrieval of episodic and semantic information. Some studies have provided evidence for a direct link between divergent thinking and episodic simulation. In two prior studies (Addis, Pan, Musicaro, & Schacter, 2016; Thakral, Yang, Addis, & Schacter, 2021), divergent thinking (as assessed in the AUT) was uniquely correlated with the number of episodic details participants generated when imagining future events, but not when remembering past events. This relationship was assumed to reflect the helpful and unconstrained nature of future thinking that can facilitate divergent thinking. In addition to this correlational evidence, when directly priming episodic retrieval via an episodic specificity induction, performance in the AUT and future simulation are enhanced (for a review, see Schacter & Madore, 2016). Moreover, neuroimaging and neuromodulation studies have revealed common and necessary hippocampal engagement during both episodic simulation and divergent thinking (e.g., Beaty et al., 2018; Thakral, Madore, Kalinowski, & Schacter, 2020; see also, Madore, Szpunar, Addis & Schacter, 2016; Madore, Thakral, Beaty, Addis, & Schacter, 2019), to suggest a common reliance of hippocampally-mediated episodic retrieval. With respect to patient evidence, semantic dementia patients show impairments in the AUT (Paulin, Roquet, Kenett, Savage, & Irish, 2020) and during episodic simulation (see above), with analogous patterns observed in patients with hippocampal damage (e.g., Duff et al., 2013; Race, Keane, Verfaellie, 2011).

## The Current Study

Considering the literature above, there appears clear evidence that episodic and semantic retrieval are involved in both divergent thinking and episodic simulation. However, both divergent thinking and episodic simulation yield unique outcomes; episodic simulation entails the creation of a first-person, novel, and future-oriented autobiographical event, whereas divergent thinking entails the generation of a creative solution to an open-ended problem. Therefore, episodic simulation and divergent thinking may differentially draw on episodic and semantic retrieval. For example, relative to divergent thinking, episodic simulation may place greater demands on episodic compared with semantic retrieval, because episodic simulation, akin to episodic memory, requires the construction of an autobiographical event. Importantly, no study to date has examined these four abilities in the same individuals, and therefore this critical question remains unanswered.

The aim of the current study is to directly test whether episodic and semantic memory retrieval uniquely relate to divergent thinking and episodic simulation. To this end, participants completed four tasks in response to object word cues: they generated meanings and definitions (semantic memory), remembered a past event (episodic memory), imagined a novel future event (episodic simulation), or generated novel uses (divergent thinking). Following prior work (e.g., Thakral et al., 2021; Addis et al., 2016; Madore & Schacter, 2016), retrieval in the episodic and semantic tasks were operationalized via the Adapted Autobiographical Interview and its ‘internal/episodic detail’ metric, a quantitative metric of the amount of retrieved information, and analogously for the episodic simulation task as a measure of the quantity of future thinking. The divergent thinking task was similarly scored for quantitative metrics such as fluency. We employed an individual differences approach using hierarchical multiple regression analyses to assess whether episodic simulation and divergent thinking performance could be predicted by performance on the episodic and semantic memory tasks.

## Specific Aims

Our first aim was to replicate prior findings where divergent thinking would be predicted by performance in the episodic simulation task, but not the episodic memory task (Addis et al., 2016; Thakral et al., 2021). To test this aim, the number of episodic details generated during episodic simulation was entered as a predictor for divergent thinking in Step 1 of the regression analysis, and the number of episodic details retrieved during episodic memory was entered as an additional predictor in Step 2 (which we predicted to be non-significant). Our second aim was to assess whether the relationship between divergent thinking and episodic simulation would hold after accounting for semantic retrieval (i.e., where episodic retrieval in Step 2 was replaced with semantic retrieval performance). In accordance with the MemiC framework (Benedek et al., 2023; see above), we predicted that divergent thinking would be predicted by both episodic simulation and semantic retrieval. Our third aim was focused on episodic simulation as the dependent variable. Here, we assessed whether episodic simulation is predicted differentially by the retrieval of episodic versus semantic information. As noted above, data from semantic dementia patients indicate that semantic memory serves as a scaffold necessary for episodic simulation. Based on these data, we hypothesized that in healthy participants, episodic simulation would be predicted by semantic but not episodic retrieval performance. However, there is also a wealth of evidence showing that future simulation and episodic memory are positively correlated across individuals (e.g., Addis, Wong, & Schacter, 2008). Therefore, episodic simulation may be predicted by both semantic and episodic retrieval. The present experimental design allows us to separately test these two possibilities.

## Material and Methods

### Participants

The experimental protocol was approved by the Institutional Review Boards of Boston College and Smith College. Written informed consent was obtained prior to participation. All participants had normal or corrected-to-normal vision, no history of neurological impairment, and were not currently taking any psychoactive medications. A total of 46 undergraduates participated in the study and received credit for a psychology course or $10/hour for participation. The final sample size consisted of 45 participants (mean age of 19.83 years [range 18-22], 23 females). One participant was excluded for not having a complete data set (e.g., only completing 3 of the 4 tasks). A power analysis using G*Power (Faul, Erdfelder, Lang, & Buchner, 2007) was conducted using the outcome of the regression analysis in the prior study of Thakral et al., (2021) predicting divergent thinking from future imagination (R^2^ = 0.16). This analysis revealed that a sample of 45 participants was sufficient for detecting a medium sized effect (power > 0.80).

### Stimuli and Task

Stimuli comprised 16 object cue words denoting common, everyday objects drawn from prior studies of memory and simulation (e.g., Madore et al., 2016, 2019; Beaty et al., 2018; Thakral et al., 2017, 2020). Cues were divided into four lists for each of four tasks (episodic memory, simulation, semantic memory, and divergent thinking). Cue lists were counterbalanced across task. An additional set of eight object cue words were used for practice items (two for each task). Trials for each task were identical with the exception of the task performed. On each trial, a cue word was initially shown at center for 500 ms, followed by a blank grey screen for 3 minutes. It was during this 3-minute grey screen that participants were instructed to perform the corresponding task (see below). Participants were free to move their eyes during the 3-minute window so long as they kept their eyes somewhere on the grey screen/monitor. After the 3-minute task period, participants were shown a series of task-specific questions and were given 5 seconds to respond to each question using the keyboard.

Each participant completed the episodic memory, episodic simulation, divergent thinking, and semantic memory tasks. Tasks were completed in blocks of four trials with task order counterbalanced across participants. Before each block, participants were instructed on the ensuing task and completed two practice trials. For each trial of the episodic memory task, participants were instructed to remember and verbally describe an event from the past few years related to the cue word. The event was instructed to be both specific in time (not lasting for more than a few minutes to a few hours) and place. They were instructed to use a first-person perspective and to verbally describe the event in as much detail as possible (including people involved, actions, and emotions) within the 3-minute time limit. After each trial of the episodic memory task, participants were asked to rate the following metrics from 1 (‘not at all’) to 5 (‘extremely’): 1) How difficult it was to perform the task, 2) How vivid/detailed the event was, 3) How personally significant the event was, and 4) How emotional the event was. Participants also indicated whether the recalled event was within the past 1-5 years (binary response: ‘yes’ or ‘no’). Verbal responses were recorded through the use of a digital voice recorder placed next to the keyboard used to respond to the tasks.

For each trial of the episodic simulation task, participants were instructed to imagine and verbally describe a novel event within the next few years related to the cue word. As in the memory task, the event was to be imagined in the first-person perspective, be specific in time and place, and be described in as much detail as possible within the 3-minute time limit. After each trial of the episodic simulation task, participants made the same ratings as in the memory task, and also rated how similar the event was to something they have experienced before (from 1 being ‘never anything similar’ to 5 being ‘this event exactly’), and how plausible the event would be if it were to occur (from 1 being ‘not at all’ to 5 being ‘extremely’). Participants also indicated whether the imagined event was within the next 1-5 years (binary response: ‘yes’ or ‘no’).

For each trial of the divergent thinking task, participants were instructed to generate as many unusual and creative uses related to the cue during the 3-minute time limit. Participants were told to be both creative and novel given past research indicating that type of instruction can impact divergent thinking (e.g., Nusbaum, Slivia, & Beaty, 2014). After each trial of the divergent thinking task, participants were asked to rate the following metrics from 1 (‘not at all’) to 5 (‘extremely’): 1) How difficult it was to perform the task, 2) How vivid/detailed the uses generated were on average, 3) How similar the uses were to experiences or thoughts they have had before, and 4) How creative (‘original and novel’) the uses were on average.

For each trial of the semantic memory task, participants were instructed to first generate two associated objects related to the cue, and then put all three objects in a sentence sorting the objects by their relative physical sizes. Once generated, participants were instructed to generate as many meanings and definitions for each object during the 3-minute time limit. These meanings and definitions could include but were not limited to typical attributes, functions, and characteristics of the objects. In contrast to the episodic tasks above, participants were instructed to generate details as if they were coming from a dictionary or encyclopedia rather than related to themselves or their own lives (i.e., to focus on meanings/definition that factually make sense given the cue and associated objects). After each trial of the semantic memory task, participants were asked to rate the following metrics from 1 (‘not at all’) to 5 (‘extremely’): 1) How difficult it was to perform the task, 2) How vivid/detailed the meaning/definitions they generated were on average, 3) How familiar they were with the objects, on average, and 4) How typical (semantically and thematically related) the objects they generated were to the cue, on average.

### Scoring and analysis

Verbal responses for each trial and task were first transcribed. Consistent with prior studies of memory and imagination (e.g., Madore & Schacter, 2016; Madore et al., 2016; Addis et al., 2008, 2016; Thakral et al., 2017, 2020, 2021), we adapted the conventional Autobiographical Interview (Levine et al., 2002) to quantify performance in each task. For the memory and simulation tasks, the central event was first identified (if more than one event was mentioned, the event discussed in greater detail was assigned as the main event interest). Once identified, the transcription was segmented into details or chunks of information (e.g., a unique occurrence or thought) and these details were categorized as internal or external following the Autobiographical Interview (Levine et al., 2002; for examples of this coding scheme, see Addis et al., 2008, 2016; Thakral et al., 2017, 2020, 2021). Internal details included the who, what, where, and when elements of the central event specific in time and place (e.g., people, actions, objects, thoughts, feelings and surroundings that focused on a central event of interest). External details included factual information, off-topic and repetitive information, commentary and metacognitive statements. For the semantic memory task, internal details were bits that contained definitions of the three words that were on-task and meaningful (e.g., for the “keys” cue an internal detail could be describing a key as a way to open a door, or that they are made of metal; see also, Madore & Schacter, 2016; Madore et al., 2016; Thakral et al., 2020). External details were bits that were off-topic, repetitive, disconnected from the definitions, or not meaningful (e.g., for the “keys” cue an external detail could be describing the task as hard, or repeating that keys are made of metal). For each participant, a mean internal and mean external detail score was computed for each event task by averaging across the 4 trials.

AUT responses were scored for standard metrics of quantitative divergent thinking (e.g., Guilford et al. 1960; Guilford 1967; Benedek et al. 2014). *Fluency* was scored as the number of uses generated, excluding repetitions. *Appropriateness* was scored as the number of uses generated that are feasible and useful in everyday life (e.g., using a using a safety pin as a laser is not appropriate). *Flexibility* was scored as the number of distinct categories that total uses can be classified under (e.g., using a safety pin as a charm for a necklace and as an earring would be classified under one category of jewelry). *Elaboration* was scored as a rating of the level of detail of each use, ranging from 0 to 2, with 0 = brief descriptions and 2 = very detailed descriptions. For each metric, the scores were averaged across the four trials. Each of the four metrics were individually z-scored, and then averaged to compute a mean divergent thinking measure (see also, Addis et al., 2016; Thakral et al., 2021). We did not score for qualitative divergent thinking such as originality/creativity, because our prior work has indicated that episodic memory and simulation are linked to *only* quantitative metrics of divergent thinking (e.g., number of uses generated) but not qualitative metrics (e.g., Addis et al., 2016; Madore et al., 2016, 2019; Thakral et al., 2020, 2021). Given these consistent findings, we chose to reduce the number of multiple comparisons (and therefore the probability of a Type-I error) by including only quantitative measures of divergent thinking in our analyses. All scoring was conducted by three raters. We obtained high interrater reliability (Cronbach’s *α* > .90) across the divergent thinking measures, as well as internal and external details for the episodic memory, simulation, and semantic memory tasks.

In our first set of behavioral analyses, we analyzed data from the memory/simulation tasks as a function of the internal and external details generated for each task. In our second set of analyses, correlation and hierarchical multiple regression analyses were used to assess the ability of episodic and semantic thinking to predict divergent thinking and future simulation performance. We first attempted to replicate the results of Addis et al. (2016) and Thakral et al. (2021), by identifying a link between the amount of internal details generated when imagining the future and divergent thinking performance, with the relationship to the amount of internal details during episodic memory being non-significant and/or weaker. To test this aim, the number of episodic details generated during episodic simulation was entered as a predictor for divergent thinking in Step 1 of the regression analysis, and the number of episodic details generated during episodic memory was entered as an additional predictor in Step 2. In a novel extension of these data, we then tested whether this relationship holds when controlling for semantic memory, by testing whether divergent thinking is predicted by internal details in the semantic memory task (i.e., where the internal detail scores during episodic memory in Step 2 were replaced with those generated during semantic memory task). Lastly, we conducted analogous regression analyses to assess the ability of internal details during episodic memory and semantic memory to predict future simulation performance. As prefaced in the Introduction, these analyses were conducted to echo findings observed in patients diagnosed with semantic dementia where semantic memory impairments lead to a selective reduction in episodic simulation relative to episodic memory ability.

Before conducting regression analyses, we confirmed that there were no violations of multicollinearity (variance inflation factor (VIF) < 5). All results are considered significant at the p < 0.05 level. For non-significant results (i.e., for the ANOVA, t-tests, and *β* estimates from the regression analyses), Bayes factors were computed using JASP software (http://jasp-stats.org/; Ly et al., 2018; Wagenmakers et al., 2018) to estimate the strength of evidence favoring the null hypothesis (Jeffreys, 1961; Dienes, 2014). We report these values as BF_10_, with values < 1 indicating substantial evidence for the null hypothesis and values near 1 indicating little or no evidence in favor of either the alternative or the null hypothesis.

## Results

### Task differences

Figure 1 illustrates the mean number of internal and external details generated for the episodic memory, episodic simulation, and semantic memory task. Following prior studies examining task performance as a function of detail type (e.g., Addis et al., 2008; Gaesser et al., 2011; Madore et al., 2014; Devitt et al., 2017), a 3 x 2 ANOVA was conducted with factors Task (episodic memory, episodic simulation, and semantic memory) and Detail Type (internal and external). The ANOVA revealed a significant main effect of Detail Type (F(1, 44) = 222.93, p < 0.001, partial *η^2^* = 0.84; Internals > Externals), main effect of Task (F(2, 88) = 37.00, p < 0.001, partial *η^2^* = 0.46), and a Task by Detail Type interaction (F(2, 88) = 7.38, p = 0.001, partial *η^2^* = 0.14). Follow-up comparisons revealed that participants generated more details for the episodic memory task relative to both of the other tasks, with participants also generating more details during episodic simulation relative to the semantic memory task (ts(44) > 3.65, ps < 0.001). To decompose the Task by Detail Type interaction, separate one-way ANOVAs were conducted on the data for each detail type. A one-way ANOVA with factor Task on the internal details was not significant (F < 1; BF_10_ = 0.15), while the analogous ANOVA on external details was significant (F(2, 88) = 38.9, p < 0.001, partial *η^2^* = 0.47). Follow-up comparisons revealed that participants generated more external details for the episodic memory task relative to both of the other tasks, with participants also generating more details during episodic simulation relative to the semantic memory task (ts(44) > 2.91, ps < 0.01, ds > 0.43).

**Figure 1.**
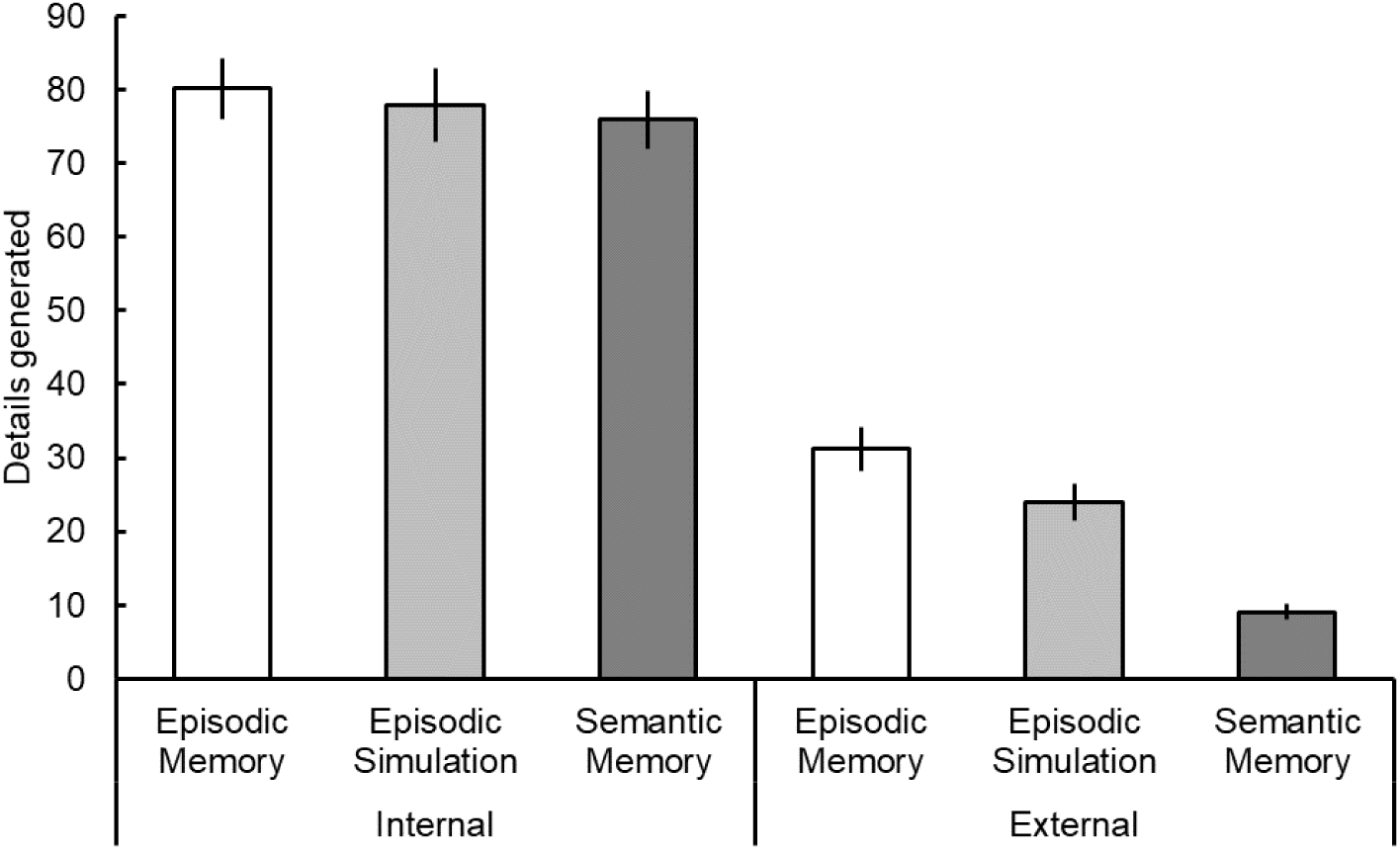
Mean number (± 1 standard error) of internal and external details generated as a function of task (episodic memory, episodic simulation, and semantic memory).

### Episodic and semantic contributions to imagining novel ideas

AUT scores were similar to those reported in our previous studies employing similar methodologies (e.g., Thakral et al., 2021, 2023; fluency mean (± 1 standard error) = 11.32 (0.60), flexibility = 7.49 (0.41), appropriateness = 11.31 (0.60), and elaboration = 1.21 (0.05)). In our first set of analyses, we tested whether creativity in the form of divergent thinking performance was correlated with the mean number of internal details comprising each type of episodic autobiographical event (remembered past and imagined future events; see Table 1 for results), and indeed it was.

**Table 1.**

|  | Episodic Memory | Episodic Simulation | Semantic Memory | Divergent Thinking |
| --- | --- | --- | --- | --- |
| Episodic Memory | 1.00 | 0.79*** | 0.69*** | 0.48*** |
| Episodic Simulation |  | 1.00 | 0.79*** | 0.49** |
| Semantic Memory |  |  | 1.00 | 0.60*** |
| Divergent Thinking |  |  |  | 1.00 |

Bivariate correlations between the mean number of internal details for the episodic memory, episodic simulation, and semantic memory tasks and divergent thinking performance (\*\*\* $p < 0.001$ , \*\* $p < 0.01$ , \* $p < 0.05$ ).

We conducted a hierarchical multiple regression where the internal details during episodic simulation was entered as a predictor for divergent thinking performance in Step 1, and the internal details during episodic memory was entered as an additional predictor in Step 2. This regression analysis (Table 2) revealed that the number of details during future simulation accounted for 24.1% of the variance in divergent thinking (*F*(1, 43) = 13.63, *p* < 0.001). The addition of the number of internal details during episodic memory in Step 2 resulted in a non-significant increase in explained variance (2.2%; *F*(1, 42) = 1.27, p > 0.20), and the overall model remained significant (*F*(2, 42) = 13.63, *p* < 0.01). Therefore, replicating our prior findings (Thakral et al., 2021) and those of Addis et al. (2016), we found that internal details comprising recalled past events did not explain a significant amount of variance in divergent thinking performance (*β* = 0.21, *t*(43) = 1.13, p > 0.20; BF_10_ = 0.50).

**Table 2.**

| **Step** | **B** | **SE B** | **$\beta$** |
| --- | --- | --- | --- |
| **1** |  |  |  |
| Constant | -0.88 | 0.26 |  |
| Internal details (episodic simulation) | 0.01 | 0.003 | 0.49*** |
| **2** |  |  |  |
| Constant | -1.08 | 0.31 |  |
| Internal details (episodic simulation) | 0.01 | 0.005 | 0.30 |
| Internal details (episodic memory) | 0.007 | 0.006 | 0.24 |

Summary of the hierarchical multiple regression analysis for internal details for the episodic simulation and episodic memory tasks in relation to divergent thinking (\*\*\* $p < 0.001$ , \*\* $p < 0.01$ , \* $p < 0.05$ ). SE, standard error.

However, imagined future events was no longer a significant predictor in the regression model with both predictors (*β* = 0.30, (*t*(43) = 1.87, *p* > 0.10; BF_10_ = 0.65).

Turning to semantic memory, number of details generated in the semantic memory task was significantly correlated with divergent thinking performance (Table 1). When the internal detail scores during episodic memory in Step 2 were replaced with those generated during semantic memory task (Table 3), there was a significant 11.7% increase in the explained variance (ΔF(1, 42) = 7.63, p < 0.01). In contrast to episodic memory, semantic memory was a significant predictor of divergent thinking (*β* = 0.56, *t*(43) = 2.76, *p* < 0.01). Once the number of internal details during semantic memory were entered as a predictor, internal details during episodic simulation was no longer a significant predictor of divergent thinking (β = 0.05, t(43) < 1; BF_10_ = 0.25).

**Table 3.**

| **Step** | ***B*** | ***SE B*** | **$\beta$** |
| --- | --- | --- | --- |
| **1** |  |  |  |
| Constant | -0.88 | 0.26 |  |
| Internal details (episodic simulation) | 0.01 | 0.003 | 0.49*** |
| **2** |  |  |  |
| Constant | -1.33 | 0.29 |  |
| Internal details (episodic simulation) | 0.001 | 0.005 | 0.05 |
| Internal details (semantic memory) | 0.02 | 0.006 | 0.56** |

Summary of the hierarchical multiple regression analysis for internal details for the episodic simulation and semantic memory tasks in relation to divergent thinking (\*\*\* $p < 0.001$ , \*\* $p < 0.01$ , \* $p < 0.05$ ). SE, standard error.

In a final analysis, we directly compared the predictive power of episodic and semantic retrieval on divergent thinking. Internal details during semantic memory was entered as a predictor for divergent thinking performance in Step 1, and the internal details during episodic memory was entered as an additional predictor in Step 2. This regression analysis (Table 4) revealed that semantic retrieval accounted for 36% of the variance in divergent thinking (*F*(1, 43) = 23.8, *p <* 0.001). The addition of the number of internal details during episodic memory in Step 2 resulted in a non-significant increase in explained variance (0.91%; *F*(1, 42) < 1). When directly comparing episodic and semantic retrieval, divergent thinking was predicted by semantic retrieval (*β* = 0.51, *t*(43) = 3.00, *p* < 0.01) and not episodic retrieval (β = 0.13, t(43) < 1; BF_10_ = 0.31).

**Table 4.**

| **Step** | **B** | **SE B** | **$\beta$** |
| --- | --- | --- | --- |
| **1** |  |  |  |
| Constant | -1.32 | 0.29 |  |
| Internal details (semantic memory) | 0.02 | 0.004 | 0.60*** |
| **2** |  |  |  |
| Constant | -1.41 | 0.31 |  |
| Internal details (semantic memory) | 0.01 | 0.005 | 0.51** |
| Internal details (episodic memory) | 0.004 | 0.005 | 0.13 |

Summary of the hierarchical multiple regression analysis for internal details for the episodic memory and semantic memory tasks in relation to divergent thinking (\*\*\* $p < 0.001$ , \*\* $p < 0.01$ , \* $p < 0.05$ ). SE, standard error.

### Episodic and semantic contributions to imagining novel future events

We next tested whether simulated future events were correlated with the number of internal details comprising remembered past events and semantic information. While both semantic memory and episodic memory were significantly correlated with episodic simulation, the correlation was numerically larger in strength for semantic memory relative to episodic memory (0.79 versus 0.69). We conducted a hierarchical multiple regression where the internal details during episodic memory was entered as a predictor for episodic simulation in Step 1, and the internal details during semantic memory was entered as an additional predictor in Step 2. This regression analysis (Table 5) revealed that internal details during episodic memory accounted for 62.7% of the variance in episodic simulation (*F*(1, 43) = 72.2, *p* < 0.001). The addition of the number of internal details during semantic memory in Step 2 resulted in a significant increase in explained variance (11.4%; ΔF(1, 42) = 18.4, p < 0.001). In contrast to imagining creative ideas (i.e., divergent thinking; Table 5), imagining future events (i.e., episodic simulation) was predicted by both episodic *and* semantic memory (*βs* > 0.0.46, *t*(43) > 4.29, *ps* < 0.001; Table 5).

**Table 5.**

| **Step** | ***B*** | ***SE B*** | **$\beta$** |
| --- | --- | --- | --- |
| **1** |  |  |  |
| Constant | 0.61 | 9.62 |  |
| Internal details (episodic memory) | 0.97 | 0.11 | 0.79*** |
| **2** |  |  |  |
| Constant | -12.88 | 8.70 |  |
| Internal details (episodic memory) | 0.58 | 0.13 | 0.47*** |
| Internal details (semantic memory) | 0.59 | 0.14 | 0.46*** |

Summary of the hierarchical multiple regression analysis for internal details for the episodic memory and semantic memory tasks in relation to episodic simulation (\*\*\* $p < 0.001$ , \*\* $p < 0.01$ , \* $p < 0.05$ ). SE, standard error.

The above analyses suggest that imagining novel future events is predicted by the ability to both remember episodic past events and retrieve semantic information. However, it is unknown whether semantic memory predicts all forms of autobiographical episodic events (i.e., those remembered and imagined) or only those imagined in the future. To answer this question, we ran a final two-step regression model with the internal details during episodic simulation entered as a predictor for episodic memory in Step 1 and the internal details during semantic memory as a predictor in Step 2 (Table 6). The addition of the number of internal details during semantic memory in Step 2 resulted in a non-significant increase in explained variance (1.03%; ΔF(1, 42) = 1.19, p > 0.20). This analysis revealed that only internal details during episodic simulation were a significant predictor of episodic memory (β = 0.66, t(43) = 4.37, p < 0.001), whereas internal details during semantic memory were not (β = 0.17, t(43) = 1.09, p > 0.20; BF_10_ = 0.26). Therefore, episodic memory is unique relative to episodic simulation: the latter was found to be exclusively predicted by semantic memory.

**Table 6.**

| **Step** | **B** | **SE B** | **$\beta$** |
| --- | --- | --- | --- |
| **1** |  |  |  |
| Constant | 29.52 | 6.48 |  |
| Internal details (episodic simulation) | 0.65 | 0.08 | 0.79*** |
| **2** |  |  |  |
| Constant | 24.83 | 7.77 |  |
| Internal details (episodic simulation) | 0.54 | 0.12 | 0.66*** |
| Internal details (semantic memory) | 0.17 | 0.16 | 0.17 |

Summary of the hierarchical multiple regression analysis for internal details for the episodic simulation and semantic memory tasks in relation to episodic memory (\*\*\* $p < 0.001$ , \*\* $p < 0.01$ , \* $p < 0.05$ ). SE, standard error.

## General Discussion

In the current study, we examined whether divergent thinking and episodic simulation are differentially predicted by episodic and semantic retrieval. In line with our initial prediction, and replicating past work (Thakral et al., 2021; Addis et al., 2016), divergent thinking was predicted by the number of episodic details generated during episodic simulation. With respect to our second aim, when directly comparing episodic and semantic retrieval, the strongest predictor of divergent thinking was semantic retrieval. Taken together, these findings align with the MemiC framework (Benedek et al., 2023). According to this framework, creative ideas generated during divergent thinking arise from four processing stages: memory search, candidate idea construction, novelty evaluation, and effectiveness evaluation. For example, when given a cue in the AUT (e.g., newspaper), individuals may search memory for objects with similar properties (e.g., notebook, umbrella) and then construct ideas based on that retrieved semantic information (e.g., use a newspaper as a makeshift umbrella). Finally, the idea is evaluated for novelty and effectiveness by running a mental simulation (e.g., based on the episodic simulation considering the effectiveness of a newspaper to protect from the MemiC framework, our findings suggest that divergent thinking, at least as assessed in the AUT, entails the interplay of both drawing information from semantic memory (e.g., bringing to mind related objects, and attributes of the cued object) and some amount of episodic simulation (e.g., running a simulation a hypothetical future event to generate candidate uses using the retrieved semantic information).

With respect to our third aim, we provide the first evidence from healthy young adults that episodic simulation is positively predicted by the retrieval of semantic information from an independent task. These findings align with the ‘semantic scaffolding hypothesis’ of episodic simulation (e.g., Irish et al., 2016). According to this hypothesis, simulating novel future events relative to remembering episodic past events relies to greater extent on semantic/conceptual information, that provides the ‘scaffold’ or framework within which simulated future events can occur and onto which relevant episodic information (e.g., who, what, where, and when information) can be integrated. A similar idea raised by Abraham and Bubic (2015) argues that future imagination emerges from elementary forms of conceptual knowledge. These proposals have been largely supported by data from semantic dementia patients, who demonstrate a selective impairment in episodic simulation but not episodic memory ability (e.g., Duval et al., 2012; Irish et al., 2012). Prior neuroimaging studies also support this view, showing greater activity in brain regions associated with semantic retrieval during episodic simulation relative to episodic memory (e.g., Addis et al., 2009, 2011; for a review, see Schacter & Addis, 2020). These findings are limited however; patient studies require interpretative caution due to the frequent occurrence of nonspecific symptomatology, and the neuroimaging studies did not directly measure semantic retrieval ability.

As expected, and replicating prior findings (e.g., Addis et al., 2008), episodic simulation performance was also positively predicted by episodic retrieval ability. Notably, the predictive power of semantic retrieval was similar to that of episodic retrieval (beta estimates of 0.47 and 0.46, respectively). In contrast to prior perspectives that have largely focused on access to episodic memory as a critical component to episodic simulation (e.g., Schacter & Addis, 2007, 2020; Sheldon & Levine, 2016), our findings suggest that such theories should be modified to equally emphasize the role of semantic retrieval, with it playing a similar role in supporting the imagination of novel future events. Relevant to this point, an important avenue for future studies is to assess whether the relative contribution of episodic and semantic retrieval may differ depending on the type of future event constructed (see, Szpunar et al., 2014). One possibility is that future events that share greater similarity to remembered past events (e.g., recast future events, see Thakral et al., 2021) may draw more on episodic relative to semantic retrieval. In contrast, when future events are novel, as those investigated here, the contribution of semantic relative to episodic retrieval may be greater. The linkage between episodic simulation and semantic memory is also particularly interesting in light of the fact that novel future events were not associated with a greater number of external details (which comprises particularly semantic information; Levine et al., 2002; see also, Renoult et al., 2020). The opposite was observed, with more external details generated during episodic memory relative to simulation (t(44) = 2.91, p < 0.01, d = 0.43). These findings suggest that the contribution of semantic memory to episodic simulation does not necessarily result in more external detail production when imagining them.

With respect to divergent thinking, the present findings replicate those of Addis et al. (2016) and Thakral et al. (2021) by showing that divergent thinking is positively predicted by episodic simulation, as indexed by the number of internal details in imagined future events. In line with our prior theorizing on the role of imagining the future in divergent thinking (e.g., Thakral et al., 2021), future thinking is particularly relevant and helpful for divergent thinking where task demands are open-ended and unconstrained, akin to future events. Importantly, our data did reveal important differences between the present and past findings. In our regression analyses, we found that while future imagination was a significant predictor of divergent thinking, it was no longer a significant predictor when both episodic simulation *and* episodic memory were included as predictors (see Table 2). In our prior studies, episodic simulation remained a significant predictor even after including episodic memory as an additional predictor of divergent thinking. A key difference between the present and past work is that in both Addis et al. (2016) and Thakral et al. (2021) participants were cued with autobiographical details (i.e., location, person and object details from the same past event) in the episodic memory task and recombined autobiographical details from different past events for the simulation tasks. Here, the stimuli were identical across the four tasks (i.e., object cues). The similarity in cues may have led to some amount of shared variance due to common cue processing across the episodic memory and simulation tasks (and therefore equally shared with the divergent thinking task). Although beyond the scope of the current study, additional work is needed to identify the specific experimental situations when divergent thinking draws on episodic retrieval. Regardless, our findings show that both divergent thinking and episodic simulation requires the retrieval of semantic information to scaffold both functions. In contrast, episodic simulation places greater demands on episodic retrieval to support the creation of an imagined future event.

In a novel extension of prior studies, we tested how semantic retrieval and future imagination ability differentially contribute to divergent thinking. Divergent thinking was predicted to a greater extent by semantic memory relative to episodic future thinking (i.e., while divergent thinking was predicted by episodic simulation ability, when considering the results of the simultaneous regression with both predictors, semantic memory was the only significant predictor of divergent thinking, Table 3). These findings align with the idea that retrieval of semantic information is critical for both the generation of novel future events (episodic simulation) and creative uses for everyday objects (divergent thinking; e.g., Roberts & Addis, 2018; Abraham & Bubic, 2015).

While our findings show that both divergent thinking and episodic simulation requires the retrieval and scaffolding of semantic information (cf., Paulin et al., 2020; Irish, & Piguet, 2013), our data also reveal important differences between these two functions. In our regression analyses, we found that while future imagination was predicted by the number of internal details participants recalled during episodic memory, divergent thinking was not (compare Tables 4 and 5). Relative to divergent thinking, episodic simulation likely requires a greater amount of event construction processes that necessitate the retrieval of autobiographical episodic details (e.g., detailed who, what, where, and when information), placing greater demands on episodic retrieval to support the creation of an imagined future event.

### Limitations

The current study focused on assessing how semantic and episodic retrieval make unique contributions to divergent thinking and episodic simulation. One important avenue for future work will be to examine how other processes might also differentially contribute to episodic simulation and divergent thinking, namely executive and attentional functions (e.g., D’Argembeau et al., 2010). Our findings show that episodic and semantic retrieval play a role in providing the building blocks for an imagined future event or novel creative idea. Also needed are executive and attentional processes that can guide and constrain the creative process by allowing for sustained attention, suppression of interference, and the selection of appropriate task strategies (for reviews, see Beaty & Schacter, 2018; Benedek & Fink, 2019; Benedek & Jauk, 2018; Roberts & Addis, 2018). As our findings clearly illustrate, episodic simulation and divergent thinking differentially draw on semantic and episodic retrieval. Therefore, there is reason to predict that each creative process might also differentially draw on executive/attentional processes.

An additional limitation stems from our operationalization of task performance. For example, we scored the divergent thinking task, the AUT, for standard quantitative metrics of performance (e.g., fluency and flexibility; for recent review, see Gerver et al., 2023). This method of scoring was chosen for two specific reasons. First, one of our primary aims was to replicate our previously reported positive correlations between future imagination and divergent thinking, and therefore we needed to adopt the same scoring method (Addis et al., 2016, Thakral et al., 2021). The fact that we replicated our findings across independent raters (not including the authors) and samples suggests that our scoring method is reliable. Nevertheless, no scoring method is perfect, and our results are limited in that respect. It will be important for future research to assess whether the current links between divergent thinking and future imagination are specific to subjective human scoring methods of the AUT or extend to automated methods that utilize large language models (e.g., Luchini et al., 2025; Organisciak, Acar, Dumas, & Berthiaume, 2023).

A similar limitation can be extended to our scoring of the episodic memory, imagination, and semantic memory tasks that were based on the adapted AI (e.g., Addis et al., 2008; Miloyan, McFarlane, Vasquez-Echeverria, 2019; Levine et al., 2002). The AI approach is widely considered as the gold standard for quantifying episodic retrieval with high reliability and convergent validity (for detailed analysis and review, see Lockrow et al., 2024). Although our study focused on quantitative measures which are assumed to reflect retrieval-related processing, use of alternative analyses may be informative to investigate other possible linkages across the cognitive functions explored here (e.g., inhibition or narrative construction; for a discussion of related issues concerning the AI, see Gaesser, Sacchetti, Addis, & Schacter, 2011; Devitt, Addis, & Schacter, 2017). In a similar vein, we examined cognitive links between divergent thinking and episodic simulation, as defined as the mental construction of a novel and future-oriented event. Our conceptualization of episodic simulation stems from a long line of theoretical and experimental work exploring the same operational definition (for reviews, see Schacter et al., 2008, 2012; Schacter & Thakral, 2023). However, episodic simulation is only one of many forms of future-oriented events (e.g., recasted future events which are not wholly novel; Thakral et al., 2021; cf., Szpunar et al., 2014). It will be important for future research to explore the relationship between divergent thinking and other forms of future-oriented cognition.

Lastly, although our results are in line with the MemiC framework (Benedek at al., 2023), we admit that our experimental design does not allow us to claim any kind of directionality (i.e., if episodic simulation is a component process of divergent thinking or vice versa). Related to this issue is whether episodic simulation and divergent thinking are in fact mutually exclusive cognitive constructs. While the MemiC framework (Benedek at al., 2023) considers episodic simulation as a component process of divergent thinking, others suggest that future imagination and divergent thinking are ‘common modes of processing’ (Roberts & Addis, 2018). Divergent thinking is classically defined as the generation of a creative solution (i.e., novel and appropriate) to an open-ended problem (Guilford, 1967). Similar to divergent thinking, episodic simulation is also inherently open-ended. Importantly, these cognitive functions are distinct in many aspects (e.g., the latter requires the generation of an autobiographical event with who, what, when and where information). Of direct relevance to this issue, we show that episodic simulation only accounted for a total of 24% of the variance in divergent thinking, and notably was no longer significant when accounting for semantic retrieval. Taken together, these findings suggest that episodic simulation and divergent thinking are in fact mutually exclusive, although further testing on this point is needed.

### Conclusion

To conclude, we demonstrate that individual differences in episodic and semantic retrieval differentially contribute to divergent thinking and episodic simulation. While episodic and semantic retrieval are important ingredients to episodic simulation, divergent thinking weighed more towards semantic relative to episodic retrieval. Therefore, findings indicate that different retrieval processes (episodic and semantic) are used to imagine novel future events and generate creative ideas.

## Data availability statement

Data are available from the first author upon request.

## Acknowledgements

We thank Tyne Burke, Keyla Cabrera, Cara Leahy, and Esther Samuel for assistance with data collection, audio transcription, and data analysis.

